# Environmental DNA reveals hidden eukaryotic diversity and fine-scale community patterns across seascape areas in the Northern Red Sea

**DOI:** 10.64898/2026.02.05.704132

**Authors:** Eva Aylagas, Karla Gonzalez, Warren R. Francis, Basmah Alabdulaziz, Joao Gabriel Rosado, Gloria Gil-Ramos, Matthew David Tietbohl Talas, Morgan Bennett-Smith, Viktor N. Peinemann, Flor Torres, Ameer A. Eweida, Michael L. Berumen, Maggie D. Johnson, Susana Carvalho

**Affiliations:** Biological and Environmental Sciences and Engineering Division (BESE), King Abdullah University of Science and Technology (KAUST), Thuwal 23955-6900, Saudi Arabia; Applied Genomic Technologies Institute, King Abdulaziz City for Science and Technology, Riyadh; Department of Biology, Boston University, 5 Cummington Mall, Boston, MA, USA, 02215; Education, Research, and Innovation Foundation, NEOM Base Camp, Building Number: 4758, Ocean Science and Solutions Applied Research Institute (OSSARI), NEOM, 49643, Saudi Arabia; Rosenstiel School of Marine and Atmospheric Sciences, University of Miami, USA; Marine Biodiversity, Ecosystems, and Conservation, NEOM, 49643, Saudi Arabia

**Keywords:** coral reef monitoring, environmental DNA (eDNA), seascape ecology, eukaryotic communities, benthic structure

## Abstract

Understanding how reef-associated biodiversity responds to seascape features is essential for monitoring and conserving coral reef ecosystems. Environmental DNA (eDNA) from seawater provides access to benthopelagic eukaryotic diversity but its relationship with benthic structure remains poorly understood. We conducted simultaneous assessment of benthopelagic eDNA derived from near-reef seawater and benthic photoquadrat surveys across 12 coral reef sites in the northern Red Sea, spanning three seascape regions: the Gulf of Aqaba, nearshore Northern Red Sea (NRS), and offshore NRS. We examined whether spatial patterns in benthopelagic eDNA communities were structured across regions and whether variation in benthic cover explained differences in eDNA-derived assemblages obtained from water samples. Benthopelagic eDNA revealed fine-scale spatial structuring across regions but showed non-significant whole-community correlation with benthic composition. When examined by major taxonomic groups, taxon-specific relationships emerged, with some taxa (i.e., *Micromonas* sp.) showing increasing relative abundances in reefs characterized by lower benthic complexity. While traditional photoquadrat surveys captured 72 benthic sessile taxa including dominant benthic groups (e.g. hard corals and algae) across four eukaryotic phyla, benthopelagic eDNA documented a broader range of eukaryotic diversity, including planktonic, cryptic, and low abundant taxa spanning 35 phyla. Notably, eDNA detected cryptic organisms overlooked by visual surveys, such as the giant clam *Tridacna* sp., even where present but not recorded in photoquadrats. Our results demonstrate that benthopelagic eDNA and visual surveys provide complementary perspectives on reef biodiversity. Rather than serving as a direct proxy for benthic structure, benthopelagic eDNA captures spatial and taxonomic patterns that may be overlooked by visual transects, supporting its use in seascape-scale biodiversity assessments and conservation planning efforts in dynamic and understudied reef systems.

## Introduction

Biodiversity loss is accelerating globally under the combined pressures of anthropogenic activities and natural environmental changes^1,2^, thus undermining our ability to effectively monitor and mitigate ecosystem alterations and respond promptly. Among the most vulnerable ecosystems on the planet are coral reefs that are increasingly threatened by climate change, overfishing, pollution, and coastal development^3,4^. These ecosystems, characterized by their structural complexity and high diversity^5,6^, support complex food webs, contribute to coastal protection, and support the livelihoods of millions of people worldwide^7,8^.

Conservation and monitoring efforts in coral reefs typically focus on conspicuous and charismatic groups such as stony corals and reef fishes^9-11^, which are often used as proxies for overall reef condition^12,13^. However, this approach overlooks a vast proportion of reef-associated biota, including small-bodied and cryptic organisms that comprise the majority of reef biodiversity^14,15^. Although less conspicuous, these organisms play critical roles in nutrient cycling, energy flow and ecosystem resilience, yet they remain poorly studied^16,17^.

Many of these taxa, including invertebrates, microeukaryotes and other sessile or cryptic organisms, are also highly sensitive to environmental disturbance, making them potential early indicators of ecological change^18,19^. Although there has been a growing recognition of the ecological relevance of these overlooked communities^20-24^, traditional biodiversity assessments relying on morphological identification often lack the resolution to detect such groups comprehensively. To overcome these limitations, it is essential to integrate methods capable of detecting such hidden diversity, thereby achieving a more comprehensive understanding of community composition and dynamics through space and time^25^. Environmental DNA (eDNA) metabarcoding has emerged as a rapid, replicable, sensitive, and scalable methodology for characterizing biological communities across an unprecedented taxonomic spectrum^26-28^. By analyzing genetic material present in water or sediment, this technique allows for the detection of rare, cryptic, and morphologically indistinct species^29^. It has proven particularly effective in biodiverse and dynamic ecosystems like coral reefs^26-28,30,31^, and serves as a valuable complement to visual surveys, enhancing biodiversity assessments^32-34^.

In coral reef systems, eDNA obtained from seawater provides a benthopelagic snapshot of biodiversity influenced by local and regional water masses that can resolve fine-scale differences in community structure across habitat types^26,31^. However, understanding the extent to which habitat composition influences benthopelagic patterns remains unexplored. Most applications of eDNA in marine monitoring remain decoupled from visual surveys, and only a few studies have directly compared eDNA-derived community profiles with benthic cover data from the same locations^35,36^. Those comparisons, however, have often focused on specific taxa, such as a direct comparison between coral cover and coral eDNA reads^37^, or between underwater visual fish survey and eDNA surveys^38^. Patterns across the broader eukaryotic communities have not been explored yet. This gap limits our ability to interpret eDNA data in ecological terms and undermines the development of habitat-specific bioindicators that could support reef assessment and management. These integrative approaches are important in dynamic and data-poor marine regions, where rapid, yet robust, assessments are needed to support evidence-based conservation planning.

In this study, we address this knowledge gap by conducting an integrated assessment of coral reef benthopelagic eDNA communities obtained from seawater samples collected above the substrate alongside photoquadrat benthic surveys across 12 reef sites in the Northern Red Sea and Gulf of Aqaba. The region has been divided into three subregions based on oceanographic hydrodynamics^39^, including nearshore reefs from the Gulf of Aqaba and the Northern Red Sea (NRS), and offshore reefs from the NRS, and is currently undergoing large-scale development, such as the construction of ports, marinas, and island resorts. Establishing robust biodiversity baselines in this region is critical for informing conservation strategies and supporting ongoing marine spatial planning in the Red Sea.

We combined high-resolution benthic imagery with eDNA metabarcoding of the mitochondrial COI gene from seawater samples collected in parallel to evaluate: i) whether spatial patterns in benthopelagic eDNA-derived eukaryotic communities are influenced by benthic structure; ii) taxon-specific relationships between benthopelagic eDNA and benthic categories and; iii) the complementarity of eDNA to detect keystone species typically identified through visual surveys (e.g. scleractinian corals). Our integrative approach may offer new insights into the strengths of eDNA-based monitoring and its utility as a complementary tool for biodiversity assessments across the coral reef seascape.

## Results

### Contrasting spatial patterns of benthic and eDNA-derived eukaryotic communities across seascape areas

A total of 72 benthic taxa and 12 abiotic categories (e.g. sand, rubble, bare rock) were characterized across the 12 sites of the study area (Figure 1) based on photoquadrat surveys. When grouped into categories, the most abundant benthic component overall was bare rock (36%), followed by live hard corals (25%), turf algae (11%), sand (9%), soft corals (7%), rubble (5%), and fleshy macroalgae (4%). The highest coral cover was recorded at SMJW (54%) and the lowest at MAAR (6%) (Figures 2, 3A), located in the Gulf of Aqaba. Sites JUQS (nearshore NRS) and MAAR (Gulf of Aqaba) were characterized by high sand cover (35% and 52%, respectively). Individuals of the giant clam *Tridacna maxima* were observed at UMHS and JQUS, nearshore reefs on the NRS representing ∼1 % of the benthic cover overall. Sites TINW and SMJW, both located in the Gulf of Aqaba, displayed the highest percentage of crustose coralline algae (CCA) (4%). Principal Component Analysis (PCA) of benthic cover data did not reveal clear clustering by seascape area, but rather by similarities in benthic category composition. Hard coral cover, sand, and bare rock combined explained 82% of the separation among samples, whilst the other 8 benthic categories combined explained 17.2% (Figure 3B).

**Figure 1:**
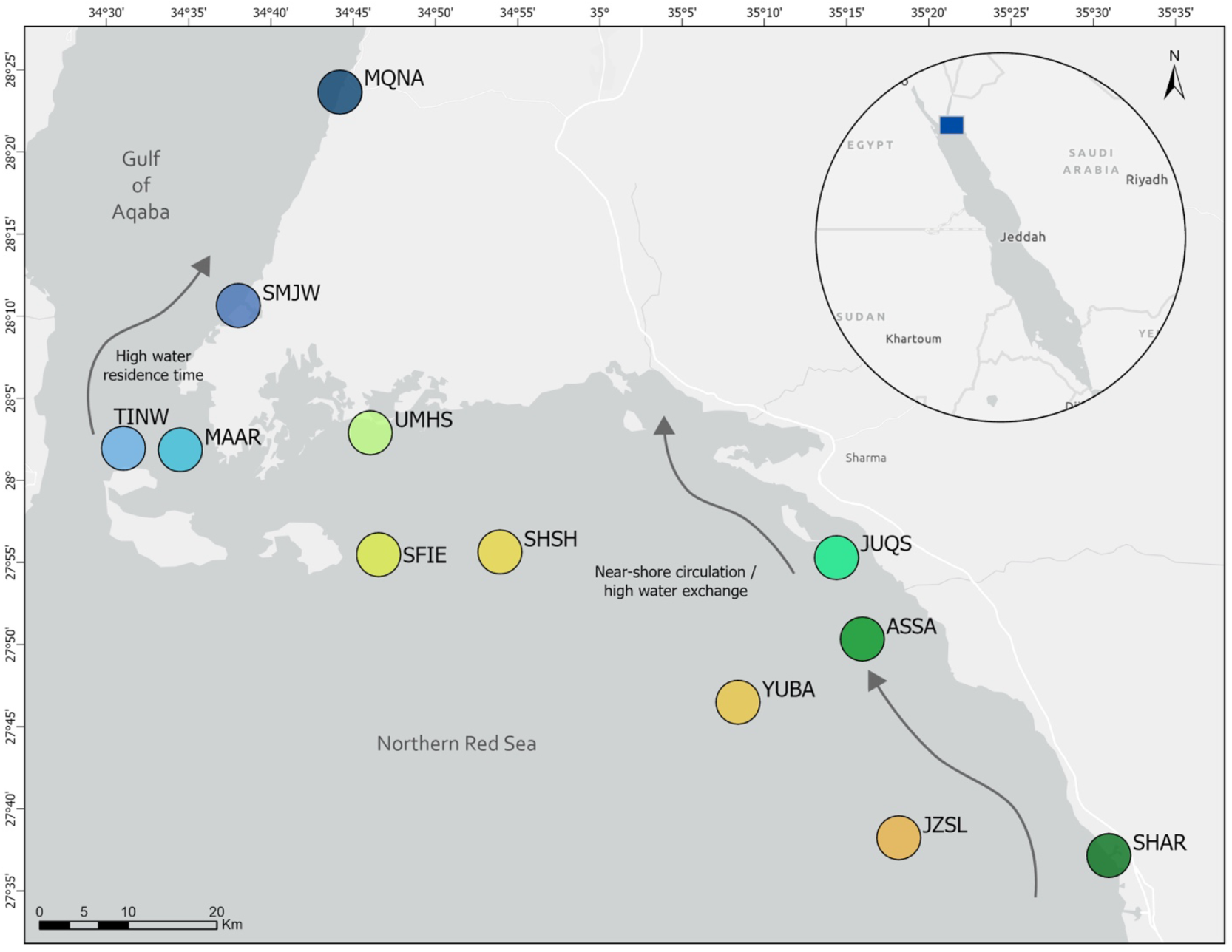
Map of the 12 sites in the Gulf of Aqaba and the Northern Red Sea. Inset shows the broader Red Sea region with the study area denoted by the blue square. Site names are: MQNA: Al Maqnah; SMJW: Sharm Mujawwan; TINW: Tiran Island NorthWest; MAAR: Marsá ‘Arīshat ar Rāshandī; UMHS: Umm Al Hassani Island; SFIE: Sanafir Island East; SHSH: Shusha Island; JUQS: Jazīrat Umm Quşūr; ASSA: Aş Şawrah; YUBA: Yob’a Island; JZSL: Jazair Sila South; SHAR: Sharm al Ḩarr. Sites are color coded based on the three seascape areas: the Gulf of Aqaba (sites in blue) nearshore Northern Red Sea (NRS) (sites in green), and offshore NRS (sites in orange), and remain consistent throughout subsequent figures. Grey arrows indicate a representation of the surface current circulation and water residence times based on Mittal et al,^39^ and Acker et al,^90^. The map was created using ArcGIS Pro software by Esri (software version 3.1.0; www.esri.com).

**Figure 2:**
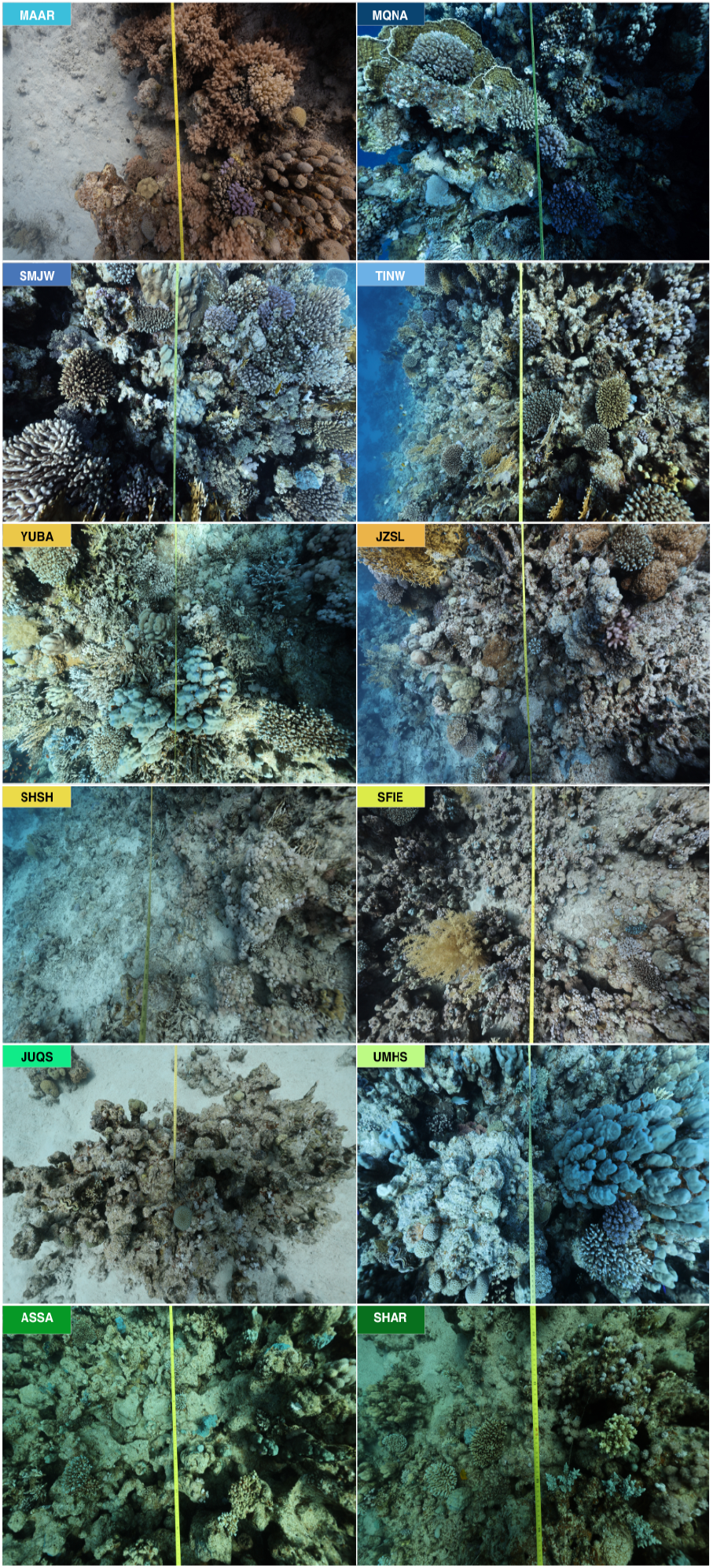
Examples of benthic photoquadrat images from each site surveyed with subregions color coded: Gulf of Aqaba - blue shades; offshore Northern Red Sea (NRS) - orange shades; inshore NRS - green shades.

**Figure 3:**
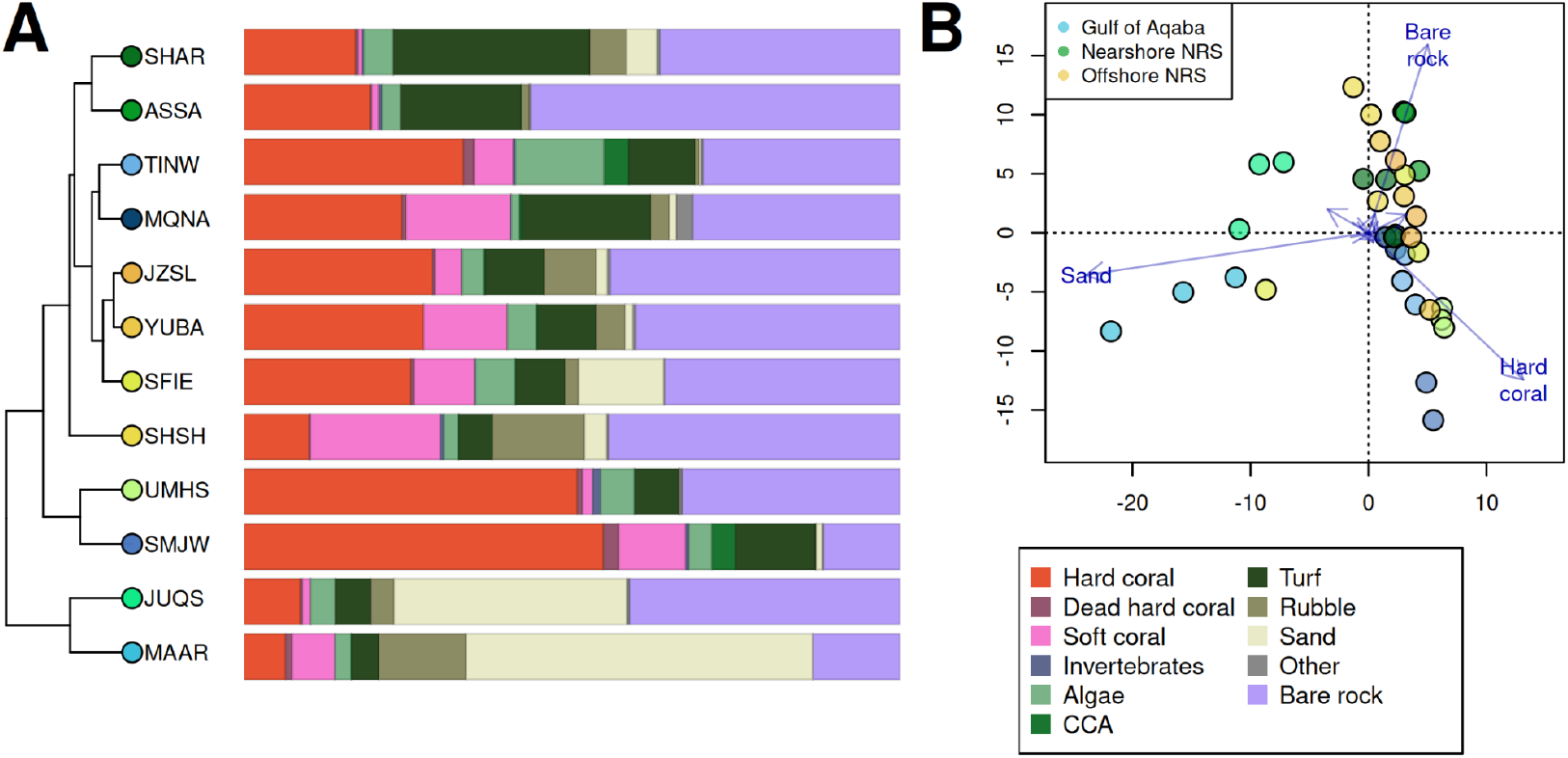
(A) Average benthic cover at the 12 surveyed sites (abbreviation as per in Fig. 1). Dendrogram displays the hierarchical clustering of sites based on the benthic cover. (B) PCA plot of the benthic cover for all transects (site SMJW is missing one transect). For clarity, only “Sand”, “Bare rock”, and “Hard coral” are shown, as the rest of the benthic categories show low contributions to the separation of the samples.

Non-metric multidimensional scaling (NMDS) based on eDNA-derived community composition revealed distinct clustering of samples (PERMANOVA, F = 8.6732, *p <* 0.001) that corresponded to the three subregions: Gulf of Aqaba, nearshore NRS and offshore NRS (Figure 4A). This spatial structuring in the benthopelagic eDNA communities by region does not fully align with the benthic habitat variation, which was instead driven by dominant benthic taxa. From the initial 8,047,190 total reads obtained from the eDNA dataset, 91.3% were retained after filtering (Supplementary Table 2), which resulted in 5,972 OTUs (97% similarity). From these, a total of 3,915 OTUs (70.7%) were assigned to phylum or lower taxonomic levels (Supplementary Figure 3), while 29.3% remained unclassified. Arthropoda was the most prevalent group across samples, representing between 18 and 60% of reads per sample (Figure 4B) followed by Cnidaria (2-39%). OTUs classified to Protists, such as algae or fungi, represented between 5.1 and 28.9% overall in terms of total number of reads. The small unicellular alga *Micromonas pusilla* (Chlorophyta) appeared as seven different OTUs and was one of the few OTUs assigned to species level that was found in all samples, accounting for 8% of total reads (varying from 2.1-22% by sample). Reads from Annelida represented between 0.005 to 10.8%, with a single replicate from SHSH that had 52% reads assigned to only a single OTU. Mollusca showed a very low diversity of OTUs across all samples (2-15 OTUs), but a significantly higher read contribution (∼18%; ANOVA, F=8.658, p=0.00451) at the nearshore reefs JUQS and UMHS, and in SMJW (Gulf of Aqaba). At these sites one OTU classified to the giant clam *Tridacna maxima* accounted for 40 - 96% of Mollusca reads, with the remaining fraction represented by other Mollusca (Figure 4C). Detection frequency of the *T. maxima* OTU across eDNA replicates was quantified to assess signal consistency. At sites where the giant clam contributed substantially to the Mollusca reads (SMJW, JUQS, UMHS), the OTU was consistently detected across replicates (SMJW: 6/6; JUQS: 5/6; UMHS: 4/6).

**Figure 4:**
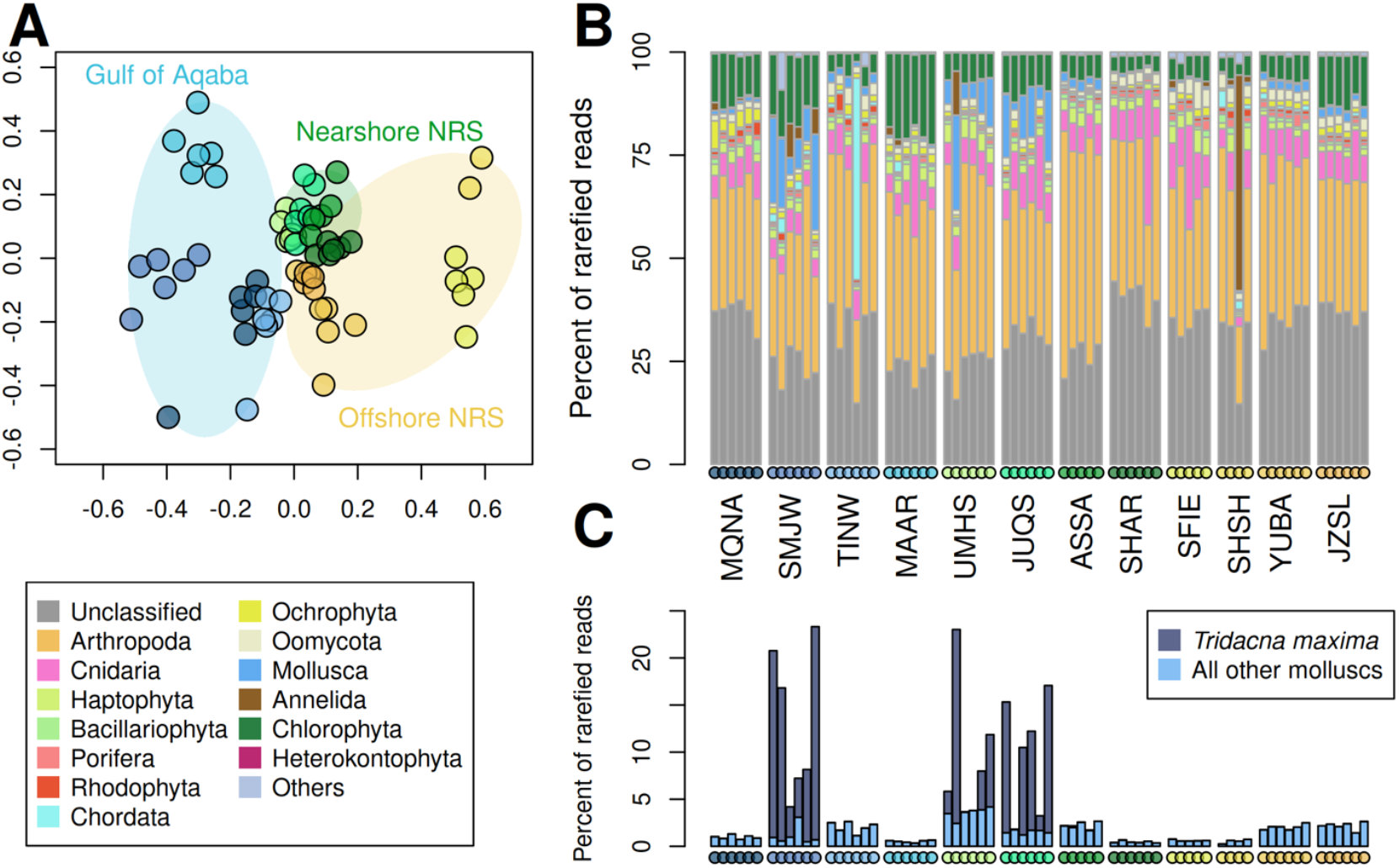
(A) NMDS plot of samples, based on Hellinger distance. Ellipses indicate subregions, with the Gulf of Aqaba in blue, nearshore NRS sites in green, and offshore NRS sites in yellow. (B) Barplot showing the relative abundance of the top 14 phyla of the eDNA dataset for each sample. All other phyla are combined into “Others”. Colored dots along the x-axis indicate the site, corresponding to the colors used in previous figures and in plot A. (C) Dominance of reads classified as *Tridacna maxima* on the total read counts at site SMJW, UMHS, and JUQS, where other molluscs account for only a few percent of reads. Order of the sites is the same as plot B.

Manual inspection of the original photographs for each transect confirmed that individuals of *T. maxima* were present at these three sites (SMJW, JUQS, UMHS), specially JUQS and UMHS. At JUQS, *T. maxima* individuals were visually detected in 26 of 60 photoquadrats (43%), whereas randomized point annotations intersected them in only 9 images (15%). At UMHS, *T. maxima* was visually present in 57 of 60 photoquadrats (95%), yet point-based annotations captured *T. maxima* in only 16 images (27%) (see Supplementary Table 4, and Supplementary Figure 4). Although eDNA data revealed the presence of the giant clam in all replicates at SMJW, only one individual of *T. maxima* was visually present in the images.

### Weak correlation between benthic structure and benthopelagic eDNA

The correlation between the dissimilarity matrices of benthic cover (e.g., hard coral, sand, algae) and the taxonomic composition of the whole eukaryotic community derived from water samples was weak and non-significant (Mantel r = –0.41, p = 0.989). Of the 72 biotic benthic categories identified from photoquadrats, 14 of the genera or families were detected in the eDNA dataset (Supplementary Figure 5). Taxon-specific relationships indicated that although whole-community eDNA dissimilarity did not correlate with benthic structure, several of the phyla displayed significant correlations with particular benthic categories (Figure 5A). For instance, Chlorophyta (dominated by *Micromonas*) showed significant negative correlations with algal cover, bare rock, and “invertebrate” cover, but a positive correlation with rubble, sand, and dead coral cover (Figure 5A-D).

**Figure 5:**
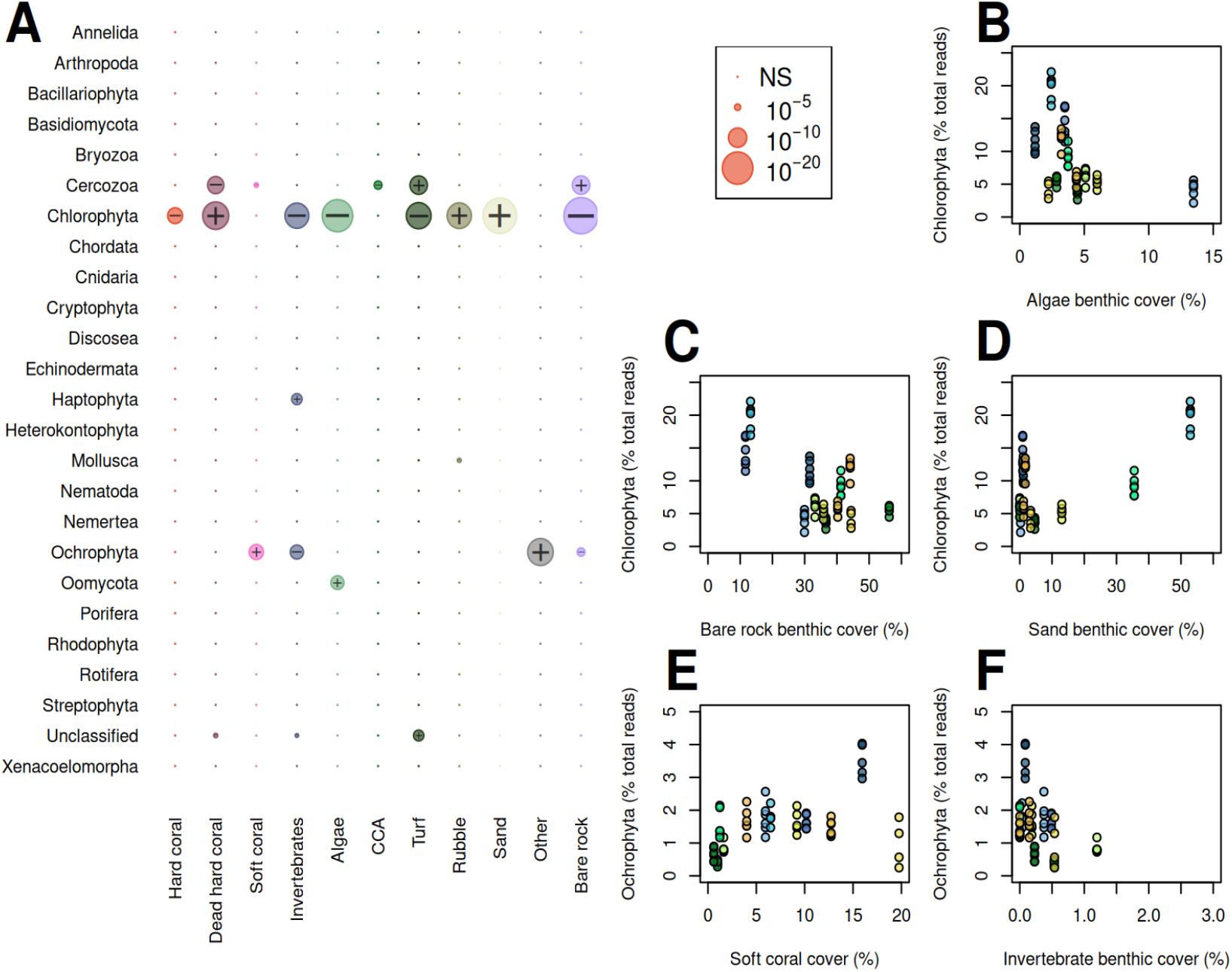
Relationship between benthic cover categories and eDNA-derived eukaryotic phyla (A) Plot of ANOVA p-values for each phylum with each category of benthic cover. Bubble size is proportional to the p-value (Bonferroni-corrected). The + and − symbols indicate the direction of the correlation, as positive and negative, respectively. (B-F) Examples of pairs with significant relationships: (B-D) Chlorophyta (Micromonas); (E-F) Ochrophyta.

While OTU relative abundance and total benthic cover of some groups revealed non-significant correlations, some groups appeared to have a similar pattern between OTU reads (%) and benthic cover. For example, the soft coral Xeniidae and the coral *Galaxea* appeared to be loosely correlated between read counts and benthic cover (Supplementary Figure 5). Such a pattern was not evident for most other benthic taxa. Several benthic categories were only detected at a single site, or others were only detected in the eDNA dataset in a single sample, such as the corals *Sarcophyton, Tubipora*, or the algae *Dictyota*, and *Liagora*. Reads of the giant clam *Tridacna maxima* were found for most samples across three sites, but at one of those sites *Tridacna* was not identified in the benthic photoquadrats.

## Discussion

Here we demonstrate that benthopelagic eDNA offers a high resolution in detecting eukaryotic community structure profiles across three coral reef subregions, uncovering biological spatial patterns that surpass those captured by traditional benthic photoquadrat surveys. By combining seawater eDNA metabarcoding analyses using the COI marker with visual assessments across 12 coral reef sites, our study confirms the capacity of benthopelagic eDNA to reveal fine-scale spatial variation in reef biodiversity^30,40^. We extend this insight by demonstrating that benthic habitat composition alone does not fully account for the observed variation in eukaryotic assemblages from the water column. The seawater samples collected directly above the reef matrix (∼50 cm off the benthos) revealed a decoupling with the benthic community observed using photoquadrats, detecting regional patterns not detected from visual transects. Notably, the biodiversity obtained from eDNA represents a combination of benthic and pelagic organisms that integrates broader biological, environmental and hydrodynamical signals, including contributions from taxa not captured within the transect belt or those hidden in the reef matrix, which are not directly tied to observed macrobenthic structure. We identified taxon-specific associations between microbial eukaryotes and particular benthic features, highlighting the potential of eDNA to uncover ecological interactions and expand our understanding of reef biodiversity beyond what is visually detected.

### Benthopelagic eDNA provides fine spatial seascape resolution and complements visually-derived benthic data

Understanding the full complexity of coral reef ecosystems requires tools capable of capturing biodiversity across spatial scales and types of organisms, including those that are microscopic, cryptic, or transient. Coral reef monitoring programs for management and conservation efforts have traditionally relied on phototransects and underwater visual census methods to characterize benthic community structure^41^. These methods are effective for documenting dominant, visually conspicuous, sessile and occasionally mobile taxa, providing essential indicators of reef condition, such as hard corals and algae. However, their limited taxonomic resolution and inability to detect cryptic, small-bodied, or morphologically indistinct organisms constrain their utility in capturing the full extent of reef biodiversity^25^. eDNA has the potential to complement these traditional approaches by revealing otherwise hidden components of the tree of life, thereby enhancing our understanding of seascape biodiversity and ecosystem complexity^28,30^. Our results demonstrate that both eDNA and phototransect methods retrieved distinct yet complementary aspects of reef-associated communities, differing in the number of taxa detected and the depth of taxonomic resolution. The photoquadrat surveys recorded 72 benthic sessile taxa across three metazoan phyla (Cnidaria, Mollusca, and Porifera) and six categories of algae. While these benthic taxa play foundational roles in reef structure and function and may be indicators of reef condition and potential community shifts^42,43^, our analysis revealed a lack of discriminatory power to distinguish among the three seascape areas assessed. This limitation has been previously reported in the central Red Sea, reinforcing the idea that traditional visual methods, due to their coarse taxonomic resolution and focus on large-sized or conspicuous organisms^44^ may limit our ability to detect ecological patterns at finer scales.

In contrast, the analysis of benthopelagic eDNA from seawater samples collected right above the reef matrix revealed 35 different eukaryotic phyla. This broader scope encompassed taxa representative from diverse range of biological assemblages including planktonic (e.g., copepods), cryptic (e.g., annelids), and microbial organisms (e.g., unicellular eukaryotes), which are either not accounted or missed in visual assessments^45^. Notably, eDNA captured groups such as Chlorophyta (99% comprised of *Micromonas pusilla*) and fungi, both of which play key roles in nutrient cycling, primary production, and symbiotic interactions^46^. These findings underscore the capacity of eDNA to provide a more comprehensive characterization of reef biodiversity, revealing critical ecosystem components that are often overlooked by traditional methods and are essential for understanding reef function and resilience at the seascape level^47^. Moreover, eDNA data showed finer spatial resolution in community composition, indicating distinct benthopelagic eukaryotic signatures across the three subregions. There has been a tendency to assume that marine eDNA is rapidly mixed in the water column and therefore unlikely to retain spatial structure or provide clear demarcation of habitats or regions^48^. However, the clear clustering of benthopelagic communities by subregion observed here suggests that seawater eDNA can retain spatially structured signals that are not always captured by traditional benthic survey techniques. The benthopelagic signal reflects a mixture of biological sources including benthic and pelagic organisms with the combined effects of eDNA decay^49^ advection and hydrodynamic mixing^50^. These physical processes are expected to vary among regions with contrasting circulation regimes, such as the Gulf of Aqaba and the Northern Red Sea^32^, and may therefore contribute to the regional structuring detected in eDNA-derived communities.

Consistent with our findings, spatial separation of eukaryotic communities derived from water samples collected close to the benthos has previously been reported on mesophotic coral reefs using samples collected ∼1 m above the seafloor^33,51^. In that study, the observed stratification was primarily driven by zooplanktonic taxa, an effect enhanced by the combined use of COI and 18S markers, which increased detection of planktonic and microbial eukaryotes. Together, these observations highlight both the strengths and limitations of seawater eDNA for reef monitoring. While benthopelagic eDNA cannot be interpreted as a direct proxy for local benthic community structure, its sensitivity to biologically and physically mediated spatial gradients makes it a powerful complementary tool for characterizing biogeographic patterns across coral reef seascapes when integrated with visual surveys.

### Bridging the gaps between eDNA and visual surveys for detection of keystone species

eDNA is transforming biodiversity detection and monitoring in coral reef ecosystems by revealing taxa often missed by traditional visual methods^52-54^ and providing higher-resolution insights into reef associated communities^30,44^. Yet, it offers a distinct and partial perspective on community structure. eDNA excels at capturing a broad spectrum of taxa but lacks the ability to provide spatial and structural information gathered from visual surveys essential for quantifying benthic structure (e.g., estimates of relative proportion of key-habitat forming taxa, reef rugosity, substrate type). As such, integrating eDNA with visual assessments has been advocated as a means for achieving comprehensive ecological knowledge of the coral reef seascape and supporting informed marine conservation planning^36^. Rather than being mutually exclusive, our findings reinforce the value of their combined use, demonstrating that integration enhances both resolution and completeness of biodiversity assessments across the coral reef seascape. Although eDNA analyses may involve additional processing time and laboratory costs, when strategically conducted (e.g. define the minimum number of samples or replicates) they can improve cost-effectiveness^18^ in addition to the gain in taxonomic breadth, spatial resolution and sensitivity (particularly for cryptic or low-abundance taxa).

In this study, we observed several informative alignments and discrepancies between eDNA and visual surveys, particularly for the giant clam *Tridacna maxima*. This species was consistently detected in benthopelagic eDNA samples at three sites (SMJW, UMHS, and JUQS). Photoquadrat analysis visually confirmed the presence of *T. maxima* at UMHS and JUQS, whilst only a single individual was recorded at SMJW using point-based annotations. Manual inspection of the original photographs revealed a higher number of individuals of *Tridacna* at UMHS and JUQS than those detected using annotation points. Yet, no additional individuals were observed at SMJW. Notably, the eDNA read proportions at SMJW were comparable to those observed at sites with visually confirmation, suggesting a strong eDNA signal of *Tridacna* despite the absence of detection in the benthic surveys. This site was characterized by high coral complexity and areas of reduced visibility, suggesting that more individuals might be hidden within the complex matrix of the reef, not captured by the photographs. The large discrepancy between visual detection rates and point-based detection of *Tridacna* highlights a methodological limitation of visual benthic surveys to quantify low-density benthic organisms, and helps explain the apparent mismatch between benthic cover estimates and consistent eDNA detection. Overall, benthopelagic eDNA detected the presence of *T. maxima* across three sites, likely capturing DNA shed into the water column via mucus, epithelial cells, or from larval stages. While additional validation using species-specific *assays*^55^ may improve detection confidence^30^, these results reinforce the value of eDNA as a complementary tool for detecting rare, cryptic or those species of conservation concern. At the same time, it highlights some detection limits inherent to traditional benthic monitoring and supports the integration of molecular tools to more comprehensively assess reef biodiversity^36^.

In coral taxa, we observed weak but consistent relationships between eDNA read relative abundance and benthic cover for several coral taxa, including *Galaxea, Goniopora*, and members of the family Xeniidae. Some genera, however, were restricted to one dataset either visual or eDNA-based (e.g., *Sarcophyton, Tubipora, Dictyota*), revealing the limitations of each method in capturing the full taxonomic spectrum. Some contrasting instances between visual and eDNA datasets were revealed by two ecologically relevant coral genera, *Pocillopora* and *Acropora*. Notably, *Pocillopora* was detected via eDNA at one site being the site with the lowest percentage cover of *Pocillopora* from photoquadrats across the 12 sites, and *Acropora* was absent from the eDNA dataset entirely. These mismatches likely reflect differences in DNA shedding rates, detection thresholds, and environmental degradation processes^33,56^. Yet, it suggests the better suitability of specific primers for the detection of corals, at least at the genus level, using eDNA^30,37,57,58^. In contrast, our study employed universal eukaryotic markers, which, while broad in scope, may compromise resolution for specific benthic groups. Whilst eDNA research moves towards optimizing its use for coral cover estimates^59^ the variability found in our study reinforces that visual surveys are better suited for estimating benthic cover and community dominance. However, eDNA is invaluable for detecting cryptic life stages, mobile organisms, or taxa residing in interstitial spaces of the reef matrix^54^.

Together, these findings emphasize that benthopelagic eDNA and visual surveys provide complementary, but not always interchangeable, insights into reef ecosystems. When combined, they improve the detection of key reef taxa, enhance ecological knowledge, and ultimately support more robust biodiversity monitoring frameworks. Yet, for direct comparisons and complementarity of benthic community assessments the use of taxa-specific markers (e.g. 28S rRNA for corals^60^), and the inclusion of additional primers for broader eukaryotic diversity (e.g. 18S rRNA) along with substrate-based sampling (e.g. sediments, scrapes of hard bottom communities) are recommended ^61^. As coral reefs face intensifying pressures, these multi-method approaches will be essential to inform adaptive conservation strategies that can support or enhance ecosystem resilience^62^.

### Benthic community composition explains limited variation in seawater eDNA profiles

Interpreting biodiversity patterns from eDNA data in an ecological context requires a clearer understanding of how these signals relate to local habitat characteristics, particularly benthic cover. While benthic habitat heterogeneity plays a fundamental role in structuring coral reef communities^63^, most eDNA-based studies have not examined the extent to which this structural variation explains patterns in the wider eDNA-derived eukaryotic community composition. Instead, most studies have focused either on generating broad inventories of reef-associated biodiversity using eDNA^36,40,45^ or on assessing the detectability of specific benthic taxa (primarily corals) through their eDNA traces in surrounding seawater^30,57,58^. Initial efforts to understand the relationship between benthic structure and the composition of crytobenthic communities (primarily invertebrates) were conducted via morphological identification of organisms collected from benthic samples, revealing strong habitat-driven community patterns^22,64,65^. In contrast, our study directly evaluated the relationship between benthic community composition and eukaryotic community dissimilarity derived from seawater eDNA, revealing no significant correlation at the whole community level.

The absence of a strong whole-community correlation suggests a partial decoupling between benthic structure and the eukaryotic assemblages detected in seawater. This likely reflects the mixed ecological origin of the benthopelagic eDNA signal captured using the COI marker gene, rather than a lack of benthic influence. Specifically, COI-based seawater eDNA integrates signals from a wide range of taxa, including both benthic-associated and planktonic organisms, with a likely overrepresentation of zooplanktonic taxa^51^. Planktonic assemblages are known to respond strongly to environmental conditions such as nutrient concentrations, turbidity, and temperature^66^,^32^, rather than directly to reef structure. These factors can therefore substantially shape planktonic communities and likely contribute to the spatial variability in benthopelagic eDNA communities observed here. Therefore, the observed non-correlation may reflect the ecological and methodological scope of the approach followed, whilst offering a broader ecological picture that integrates biological interactions and environmental conditions. This emphasizes the value of eDNA as a complementary tool in reef monitoring. However, interpretation of COI metabarcoding data is constrained by its broad taxonomic coverage and the relatively high proportion of unclassified OTUs, which limits inference to a subset of the recovered diversity^51^. Importantly, DNA metabarcoding cannot distinguish between life stages^67^. As a result, sequences taxonomically assigned to benthic organisms may represent planktonic larval stages rather than benthic adults, further complicating the interpretation of seawater eDNA in relation to benthic structure.

Addressing these limitations will likely require multi-marker approaches combined with improved and more comprehensive DNA reference databases to better resolve taxonomic identity and to clarify which components of the community are driving observed spatial patterns. In addition, integrative studies that disentangle the relative contributions of habitat structure, environmental conditions, and biological processes^32^ will be critical for resolving the complex interactions between reef fauna, planktonic assemblages, and abiotic drivers shaping eDNA community profiles.

Despite the weak correspondence at the whole-eDNA community, taxon-specific relationships between eDNA and benthic categories were detected. In particular, Chlorophyta, dominated by the picoeukaryotic green algae *Micromonas pusilla*, exhibited significant correlations with several benthic categories. Relative abundances of *Micromonas* were lower at sites with high live coral or macroalgal cover and higher at sites dominated by dead coral and rubble. Whilst, these patterns may suggest a preference towards reef habitats with reduced benthic complexity or areas undergoing degradation, this result needs to be further validated with environmental covariates and multi-maker approaches. *Micromonas* is globally distributed and has been reported as a primary producer in oligotrophic coastal and oceanic waters^68,69^ as well as coral reef ecosystems^26^. However, its functional role in reef environments is poorly understood. Our findings provide evidence that changes in the relative abundance of *Micromonas* may respond to benthic structure potentially due to increased penetration of light, or elevated nutrient availability. Recent work suggested that the prevalence of *Micromonas* may be attributed to its flexible trophic strategy relying on both photosynthesis (i.e. autotrophy) and the uptake of organic matter or bacteria (i.e. mixotrophy) for growth^68^. In this context, where visual surveys may overlook fine-scale microbial dynamics, micro-eukaryotes may offer valuable insights into habitat conditions and serve as early indicators of potential shifts in coral-associated communities or elevated nutrient input^70^. Together, these findigns highlight the capacity of benthopelagic eDNA to reveal meaningful ecological patterns^19^ and support the inclusion of microbial taxa in reef monitoring programs^71,72^. Thus, we advocate for future studies to incorporate markers more specifically targeting eukaryotic diversity (e.g. 18S rRNA), alongside COI, to better disentangle the associations between benthic and pelagic reef-associated assemblages.

## Conclusion

Our study highlights the suitability of eDNA metabarcoding to complement traditional coral reef monitoring approaches. While visual surveys provide spatially explicit data on dominant benthic taxa, eDNA offers greater taxonomic resolution and captures additional dimensions of biodiversity not accessible by visual or morphology-based methods alone. Rather than serving as a proxy for benthic habitat composition, benthopelagic eDNA derived from near-reef seawater captures fine-scale seascape signatures structured by regional hydrodynamics, while still detecting taxon-specific shedding from sessile organisms (e.g., *Tridacna maxima*). This highlights the complementarity of appraoches: benthic photoquadrat surveys quantify structural habitat and dominance, whereas eDNA expands taxonomic breadth and reveals cryptic or low-density taxa not readily captured by visual methods. In fast-developing marine regions like the Northern Red Sea, where timely, scalable, and comprehensive tools are needed to guide conservation efforts, eDNA holds promise as a critical component of future biodiversity monitoring. Its ability to resolve fine-scale spatial patterns, detect early ecological changes, and uncover hidden diversity makes it an invaluable addition to the coral reef monitoring toolbox.

## Material and Methods

### Study area

The study area is located along the northern Saudi Arabian Red Sea coast and the southern east Gulf of Aqaba (Figure 1). Surveys were conducted in June 2023 at 12 coral reef sites. Four of these sites were in the Gulf of Aqaba (Al Maqnah, MQNA (28.394, 34.736); Sharm Mujawwan, SMJW (28.177, 34.634); Tiran Island North West, TINW (28.033, 34.517); and Marsá ‘Arīshat ar Rāshandī, MAAR (28.03, 34.575)). In the Northern Red Sea (NRS), four sites were located on nearshore reefs (Umm Al Hassani Island, UMHS (28.048, 34.767); Jazīrat Umm Quşūr, JUQS (27.922, 35.24); Aş Şawrah, ASSA (27.839, 35.266); and Sharm al Ḩarr, SHAR (27.62, 35.516)) within 7 km of the coastline, and four sites were on offshore reefs (Sanafir Island East, SFIE (27.924, 34.776); Shusha Island, SHSH (27.927, 34.898); Yob’a Island, YUBA (27.775, 35.14); and Jazair Sila South, JZSL (27.637, 35.302)) at distances greater than 13.5 km from the shore. This spatial delineation into three seascape areas (Gulf of Aqaba, nearshore NRS, and offshore NRS) was made according to specific oceanographic hydrodynamics, where nearshore reefs in the NRS are more directly influenced by coastal currents, while offshore waters experience slower and more dispersed water movement^39^. Sites were therefore selected within these three distinct seascapes to generate a spatial reference for community patterns and to evaluate whether benthic and eukaryotic communities revealed by eDNA exhibit consistent spatial structuring across regions.

### Benthic community data collection

At each site, benthic surveys were conducted along triplicate 50m semi-permanent transects placed at ∼ 10 m depth, following the contour of the reef. Photographs of the benthos (1 m^2^), referred to as photoquadrats, were taken every meter along the transects (50 photoquadrats per transect) with a camera Canon EOS R5. From these, the first 20 highest-quality photographs were selected for subsequent analysis, based on focus and sharpness. Images were manually annotated in CoralNet^73^, to determine benthic coverage of taxa, with 48 random points per photo. The benthos under each point was classified into one of 83 distinct benthic biotic and abiotic categories (Supplementary Table 1), previously described in Gonzalez et al.^74^. Data were then grouped into broader categories (i.e., benthic groups), such as hard corals (including *Millepora*), soft corals, fleshy macroalgae, turf algae, crustose coralline algae (CCA), invertebrates (e.g., sponges, *Tridacna* giant clams), bare rock, rubble, dead hard coral, and sand. Organisms such as hard corals, soft corals, and macroalgae were identified to the genus level when the image resolution allowed. Additionally, colonies of *Acropora* and *Porites* were further categorized based on their growth forms (e.g., branching, massive, encrusting, table and columnar). Points falling on the transect tape and shadows were excluded from analysis, and the remaining annotations were adjusted to represent 100% of the benthic cover. The percent cover of each benthic category across all 20 photographs of each transect was averaged. Further, the average of the three replicate transects was calculated to obtain the percent cover at the site level.

### Benthopelagic eDNA-derived eukaryotic community sample collection

At each sampling site, six replicate water samples (2 L) were collected 50 cm above the substrate using Nalgene bottles. Each bottle (1 per replicate sample) was filled with MiliQ water on the surface and submerged by a diver. At the bottom, the bottle was inverted, opened and purged using air from the regulator to get the MilliQ out of the sampling bottle. The sampling bottle was immediately filled with seawater, closed and transported in a mesh bag to the surface. Following this approach, two replicate seawater samples were collected just above each triplicate 50m semi-permanent transects. Each water sample was filtered on board the vessel through one individually packaged sterile 0.22 μm (47 mm diameter) filter membrane (Millipore) placed on each adaptor holder of a 6-branch aluminum manifold Vacuum Filtration System. Negative control samples were obtained by filtering 500 mL of MilliQ water treated the same as an environmental sample. Each filter was placed into individual 5 mL cryogenic tubes and submerged in liquid nitrogen for the duration of the expedition (up to 2 weeks, depending on the site). All equipment and material in contact with water samples and filter membranes was soaked in a 10% bleach solution for 5 min and rinsed three times with MilliQ water between each sampling event to prevent cross-contamination. Once in the laboratory, the filters were stored at −80°C for further DNA extraction.

DNA extractions were conducted at approximately three weeks after collection following the protocol outlined by DiBattista et al.^75^ One third of each filter was used for DNA extraction keeping the remaining filter as a backup at −80°C. Each third was then cut into small pieces using sterile forceps and scissors and placed into a 1.5 mL sterile Eppendorf tube containing 540 μL Buffer ATL and 60 μL proteinase K. Samples were incubated at 56°C for 3 h in a ThermoMixer (350 rpm) and vortexed hourly. After incubation, the supernatant was transferred into a new Eppendorf tube and the DNA extracted using the DNeasy Blood and Tissue Kit (QIAGEN), adding 400 μL of AL buffer and 400 μL of ethanol. Blank extraction controls were included in each DNA extraction following the same protocol. The Qubit dsDNA High Sensitivity assay Kit (Invitrogen, Thermo Fisher Scientific) was used to measure DNA concentration. Each DNA sample was normalized to 5 ng/μL using nuclease-free water and stored at −20°C until further processing.

A 313 bp region of the cytochrome c oxidase subunit (COI) gene was amplified in triplicate from each DNA sample using the forward 5’ - 3’ mlCOIint (GGWACWGGWTGAACWGTWTAYCCYCC) and reverse 5’ - 3’ jgHCO2198 (TAIACYTCIGGRTGICCRAARAAYCA) universal primers^76^ containing Illumina sequencing adapters. Each PCR reaction consisted of 1 mL of 10mM of each forward and reverse primer, 12.5 mL of KAPA HiFi HotStart DNA Polymerase (HotStart and Ready Mix formulation, KAPA Biosystems), 8.5 mL of RNA-free water, and 2 mL of genomic DNA. PCR conditions consisted of an initial 3-min denaturation step at 98°C, followed by 35 cycles of 20s at 98°C, 60s at 46°C, and 90s at 72°C and finally 5 min at 72°C. A unique combination of Illumina dual indexes (Nextera™ XT Index Kit) was attached to the Illumina sequencing adapters of each pooled sample in a second PCR. DNA libraries were loaded into the Illumina MiSeq sequencing platform (v3 chemistry) at KAUST (Bioscience CORE Labs) to determine sequence reads. Twenty percent phiX was included for internal control. Sequence data were automatically demultiplexed using MiSeq Reporter (v2), and forward and reverse reads were assigned to samples.

The sequencing dataset included six replicate samples per site, with one site having four and one having five, for a total of 69 samples and 14 PCR blanks (typically one for each site), for a total of 84 sets of paired-end reads. Primer sequences from each forward and reverse read were trimmed using cutadapt^77^, with options “-e 0.07 --discard-untrimmed -g ^GGWACWGGWTGAACWGTWTAYCCYCC –G ^TAIACYTCIGGRTGICCRAARAAYCA” discarding read pairs where either the forward or reverse primer was not found. Reads were then filtered and denoised using the DADA2 pipeline within R^78^. A maximum number of “expected errors” (maxEE) of 2 were allowed for the forward reads and 4 for the reverse reads. Reads were only retained if the length was exactly 313 bp to remove possible pseudogenes^79^ or one complete codon longer (316 bp) or shorter (310 bp), which were the vast majority of ASVs (Supplementary Figure 1). The BOLDigger3 tool^80^ along with BLASTn^81^ coupled with the lcaPident function in the biohelper R package (https://github.com/olar785/biohelper/) were used to assign taxonomy to these chimera-, contaminant- and nuMT-free ASV sequences (BLASTn parameters. Specifically, for each ASV, up to 5 top BLAST hits were retrieved using a stringent e-value of 1e-10 based on the curated BOLD database v5^82^ as well as the NCBI core nucleotide (nt) database (version 5; January, 2025). Taxonomic assignment was determined using the lcaPident approach, which assigns sequences to the lowest common ancestor supported by sequence similarity across hits. Singleton ASVs or ASVs detected in <2 libraries were excluded along with non-target ASVs representing taxa outside of Eukaryota. A total of 9,161 COI ASVs were retained after the filtering steps. Denoising was followed by clustering, as recommended for highly variable mitochondrial markers such as COI^83-85^. ASVs were clustered at 97% sequence identity^86^ using VSEARCH^87^ with default parameters, resulting in 5,972 operational taxonomic units (OTUs). Counts for each OTU were summed using a custom Python script “merge_uc_clusters.py” (see https://github.com/BEMlabKAUST/NEOM-2023-eDNA-COI).

### Analysis of community patterns and composition

One eDNA sample, Site-ASSA-R6 had ∼10x fewer reads than most other samples (Supplemental Figure 2), and was excluded from further analysis, and OTUs found only in that site were removed from the dataset. The remaining 68 samples varied between 21,014 and 209,069 reads, though most had between 30,000-70,000 reads, and were then rarefied using the R function rrarefy() to generate a random subset with 20,000 reads each used for further analysis. Multivariate analyses were conducted on both the benthic cover data and the eDNA-derived taxonomic profiles to assess patterns in community composition. For the benthic dataset, Principal Component Analysis (PCA) was performed using the pca() function in the vegan package^88^. Vectors representing the benthic category contributing the highest to the separation of samples were overlaid on the ordination. For the eDNA dataset, Non-metric Multidimensional Scaling (NMDS) using the metaMDS() function in the vegan package was used to visualize patterns in eukaryotic community composition among sites based on Hellinger distance^89^. To do that, the Hellinger transformation was applied to the rarefied OTU table (20,000 reads), to reflect community composition (not absolute read counts) while reducing the dominance of highly abundant taxa without discarding relative abundance information.To statistically test for differences in community composition across subregions (Gulf of Aqaba, nearshore NRS, and offshore NRS), a Permutational Multivariate Analysis of Variance (PERMANOVA) was done using the adonis2() function in vegan (1 factor: subregion; 3 levels: Gulf of Aqaba, nearshore NRS, and offshore NRS).

To evaluate the degree of relationship between benthic categories and eDNA-derived community datasets, a Mantel test was done using the mantel() function in the vegan package, comparing Bray–Curtis dissimilarity matrices of benthic cover and eDNA community composition. Additionally, to assess whether relationships were detectable within specific components of the community, we performed group-level analyses by testing correlations between benthic cover categories and the relative abundance of major eukaryotic phyla detected in the eDNA dataset (after filtering low abundant phyla: < 100 reads), using ANOVA from the aov() function, Bonferroni-corrected for 275 tests. Validation of *Tridacna* detection in photoquadrat surveys was conducted at sites where either *Tridacna* was intercepted by randomized annotation points in at least five images, or *Tridacna* reads were detected in at least three of the six replicate eDNA samples. For each transect at these sites, all photoquadrats included in the benthic analysis were visually inspected to record: (i) the number of images in which *Tridacna* individuals were visually present; (ii) the number of *Tridacna* individuals manually detected across images (iii) the number of images in which at least one randomized annotation point used for CoralNet-based benthic cover estimation intersected *Tridacna*; and (iv) the number of *Tridacna* detections based on annotation points. The proportion of images in which *Tridacna* individuals were visually present was then compared with the proportion of images in which *Tridacna* was captured by randomized point annotations.

## Acknowledgments

We would like to express our special thanks to Chakkiath Paul Antony for his support during raw sequencing data processing and to Carolina Bocanegra and Doaa Baker from their assistance in sample preparation.

## Ethical approval and permits

Field sampling was conducted in compliance with institutional biosafety and bioethics regulations. The study was approved by the KAUST Institutional Animal Care and Use Committee under the reference #24IACUC020 and seawater samples were collected under the permits issued by NEOM.

## Author Contributions

S.C., E.A., F.T., and A.E. funded the project and contributed to study design. M.D.T.T., M.B.S., V.N.P., B.A., J.G.R., G.G.R. contributed to sample acquisition and laboratory processing. E.A., K.G., and W.R.F. analyzed the data and produced the figures. E.A., K.G., W.R.F, and S.C. wrote the manuscript. M.D.J., M.L.B., S.C., and E.A. supervised students contributing to this manuscript. All authors revised and contributed to the final version manuscript.

## Data availability

Datasets and scripts used for data analysis have been deposited in GitHub https://github.com/BEMlabKAUST/NEOM-2023-eDNA-COI

Reads have been deposited in NCBI SRA under BioProject accession PRJNA1283908.

