## Supplementary figures and tables for "Environmental DNA reveals hidden eukaryotic diversity and fine-scale community patterns across seascape areas in the Northern Red Sea"

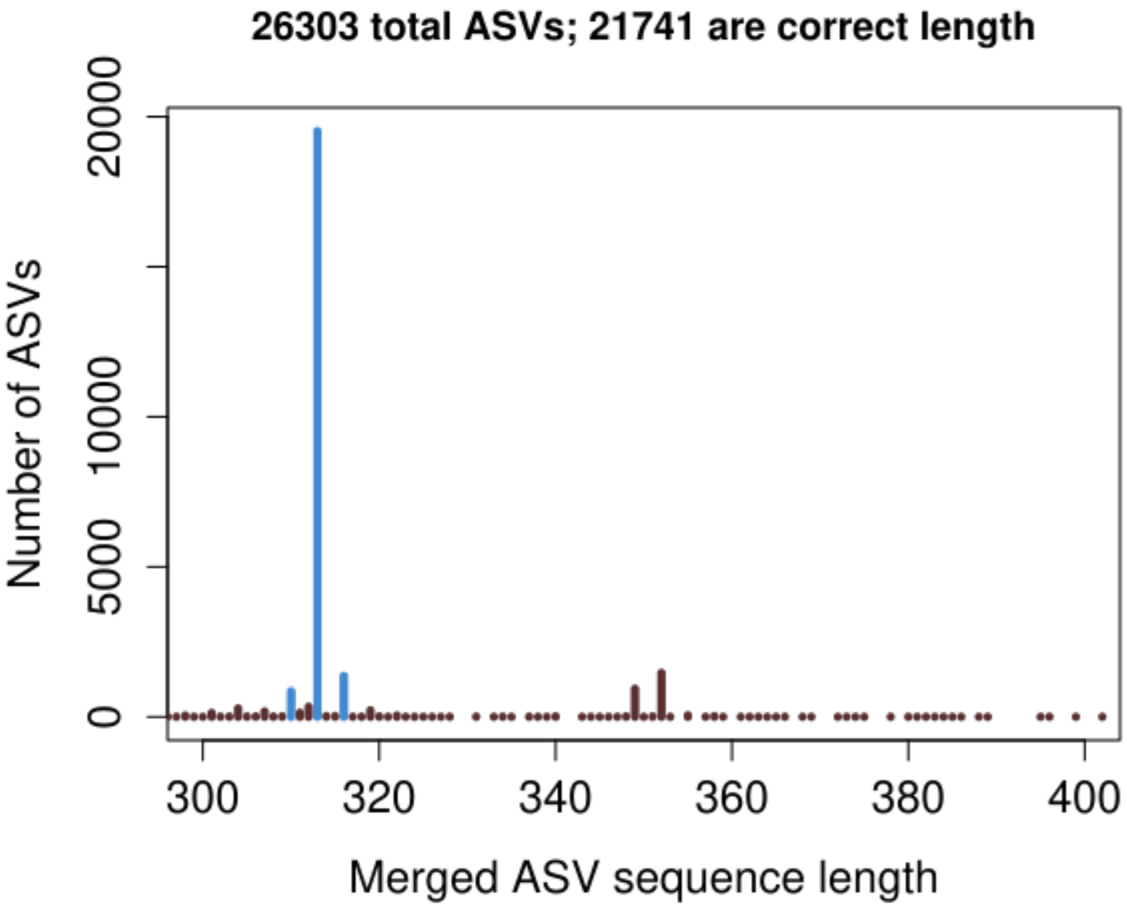


**Supplementary Figure 1.** Histogram of the length of ASVs. Only the peaks in blue (310, 313, and 316 bp) were retained for further analysis.

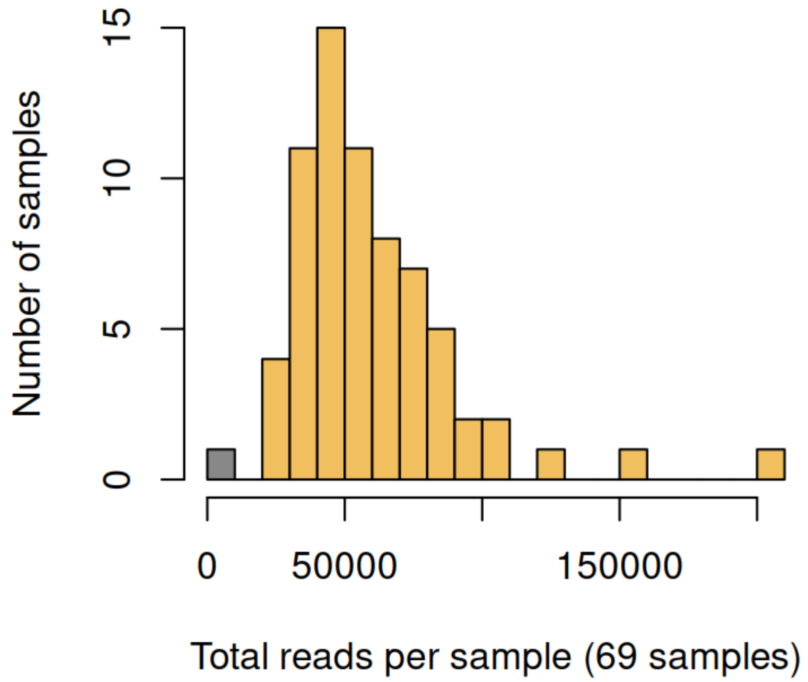


**Supplementary Figure 2:** Total number of reads for each sample. The gray sample corresponding to site ASSA-R6 was removed from subsequent analyses.

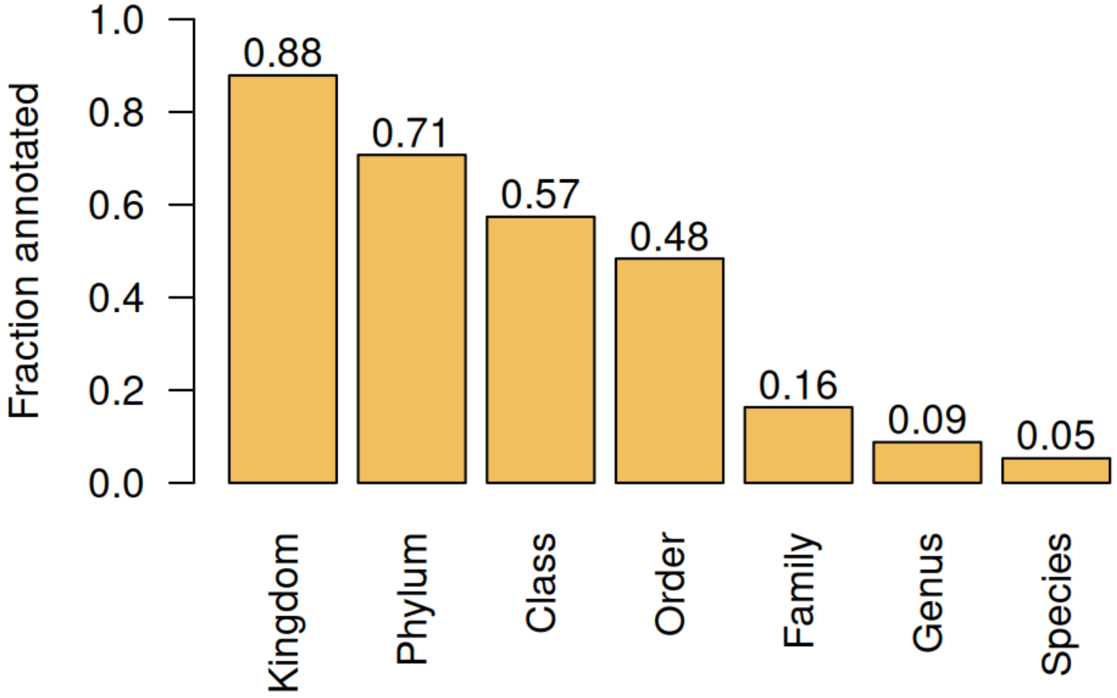


**Supplementary Figure 3:** Fraction of OTUs assigned to Kingdom, Phylum, Class, Order, Family, Genus and Species level.

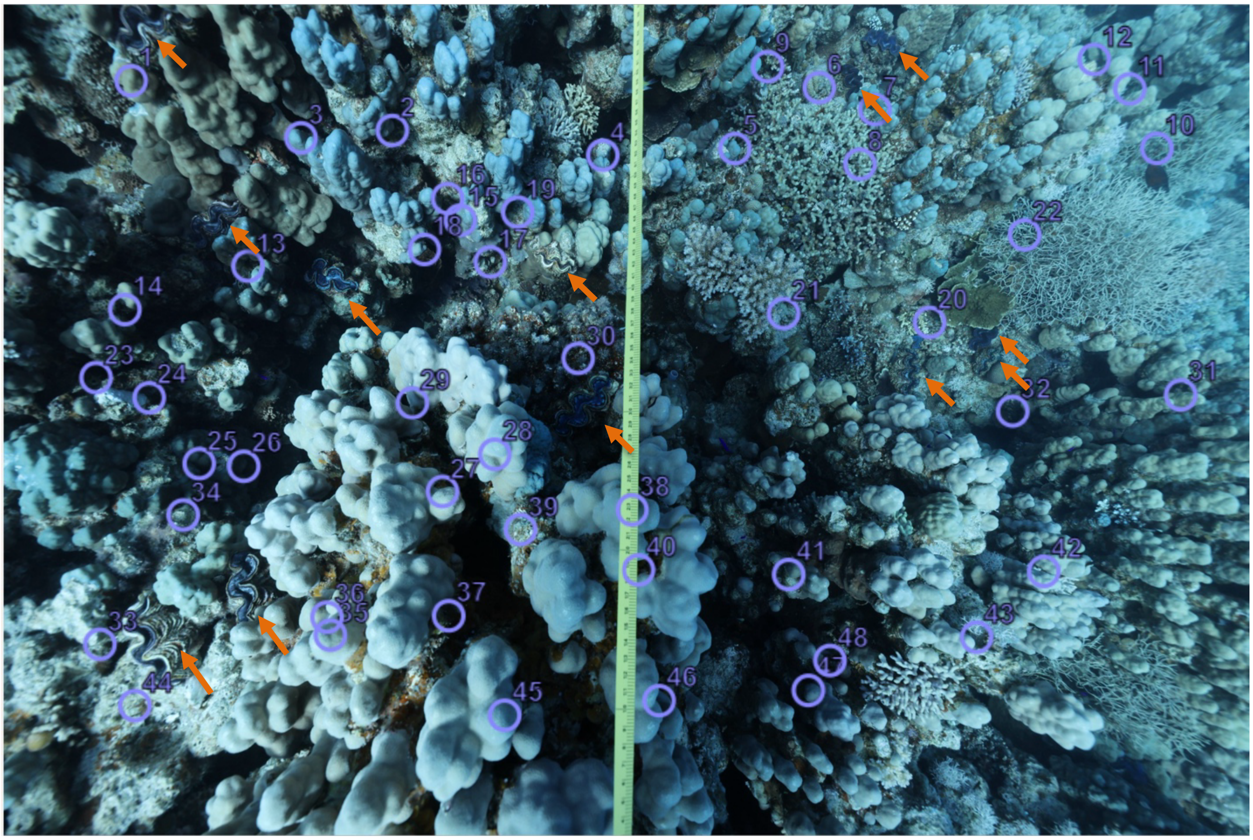

**Supplementary Figure 4.** Example photoquadrat image from reef site UMHS used for benthic analysis in CoralNet. Randomized annotation points are shown as purple circles with their corresponding identification numbers. Orange arrows indicate individuals of *Tridacna maxima* visually detected during manual inspection that fall outside the randomized point locations.

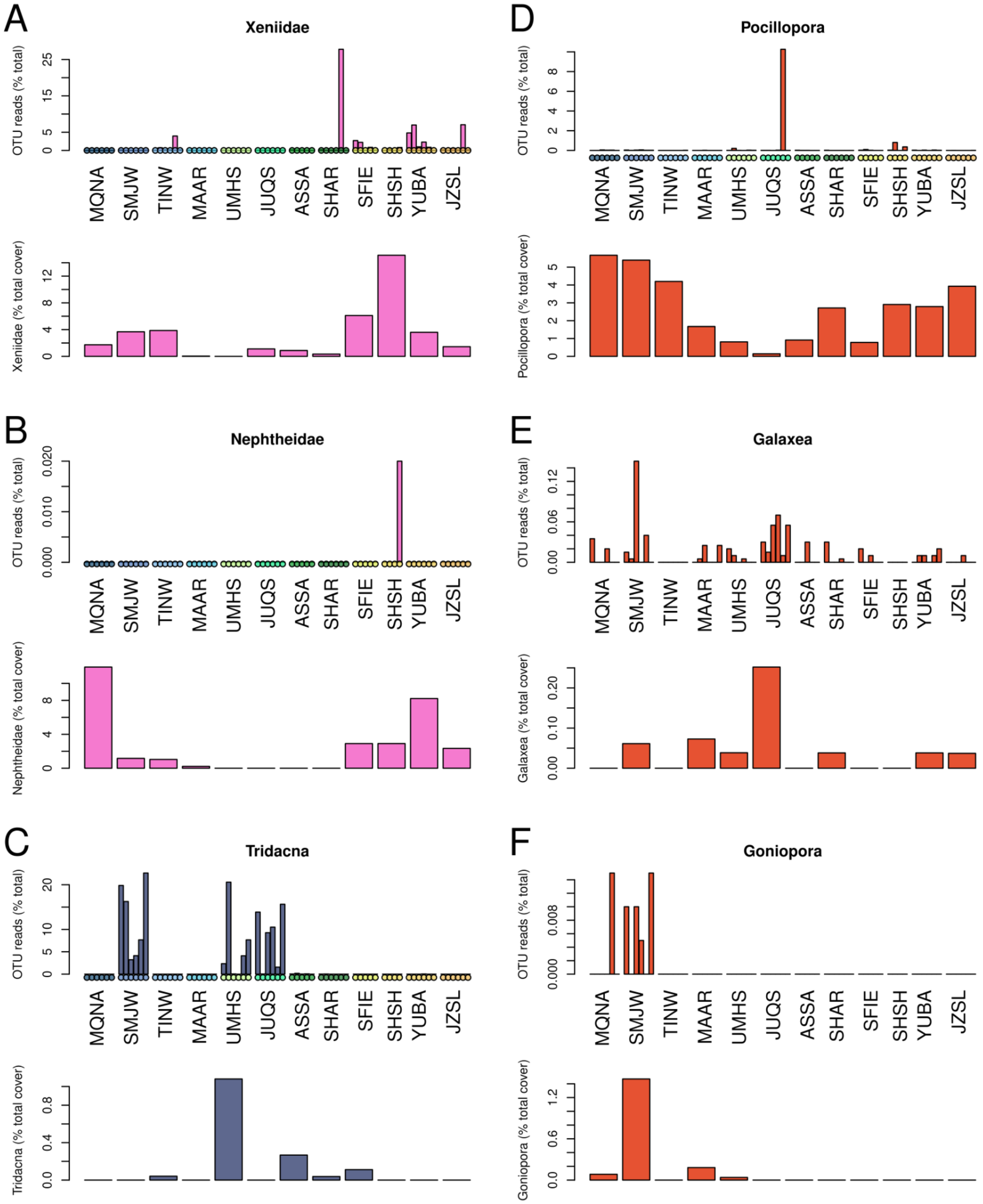
A
Xeniidae
OTU reads (% total)
Xeniidae (% total cover)
B
Nephtheidae
OTU reads (% total)
Nephtheidae (% total cover)
C
Tridacna
OTU reads (% total)
Tridacna (% total cover)
D
Pocillopora
OTU reads (% total)
Pocillopora (% total cover)
E
Galaxea
OTU reads (% total)
Galaxea (% total cover)
F
Goniopora
OTU reads (% total)
Goniopora (% total cover)
MQNA
SMJW
TINW
MAAR
UMHS
JUQS
ASSA
SHAR
SFIE
SHSH
YUBA
JZSL

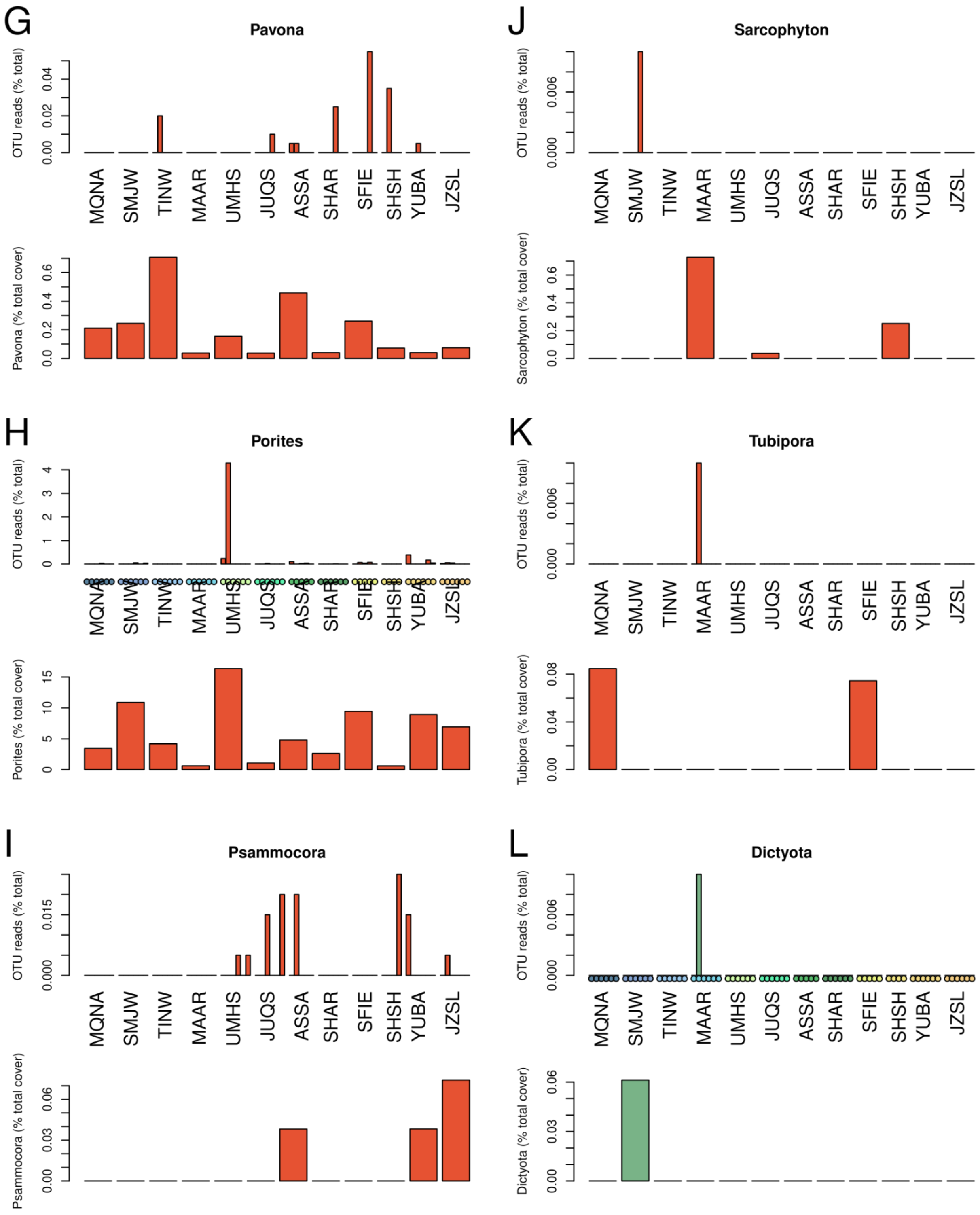

G
Pavona
OTU reads (% total)
0.00 0.02 0.04
MQNA SMJW TINW MAAR UMHS JUQS ASSA SHAR SFIE SHSH YUBA JZSL
Pavona (% total cover)
0.0 0.2 0.4 0.6
J
Sarcophyton
OTU reads (% total)
0.000 0.006
MQNA SMJW TINW MAAR UMHS JUQS ASSA SHAR SFIE SHSH YUBA JZSL
Sarcophyton (% total cover)
0.0 0.2 0.4 0.6
H
Porites
OTU reads (% total)
0 1 2 3 4
MQNA SMJW TINW MAAR UMHS JUQS ASSA SHAR SFIE SHSH YUBA JZSL
Porites (% total cover)
0 5 10 15
K
Tubipora
OTU reads (% total)
0.000 0.006
MQNA SMJW TINW MAAR UMHS JUQS ASSA SHAR SFIE SHSH YUBA JZSL
Tubipora (% total cover)
0.00 0.04 0.08
I
Psammocora
OTU reads (% total)
0.000 0.015
MQNA SMJW TINW MAAR UMHS JUQS ASSA SHAR SFIE SHSH YUBA JZSL
Psammocora (% total cover)
0.00 0.03 0.06
L
Dictyota
OTU reads (% total)
0.000 0.006
MQNA SMJW TINW MAAR UMHS JUQS ASSA SHAR SFIE SHSH YUBA JZSL
Dictyota (% total cover)
0.00 0.03 0.06

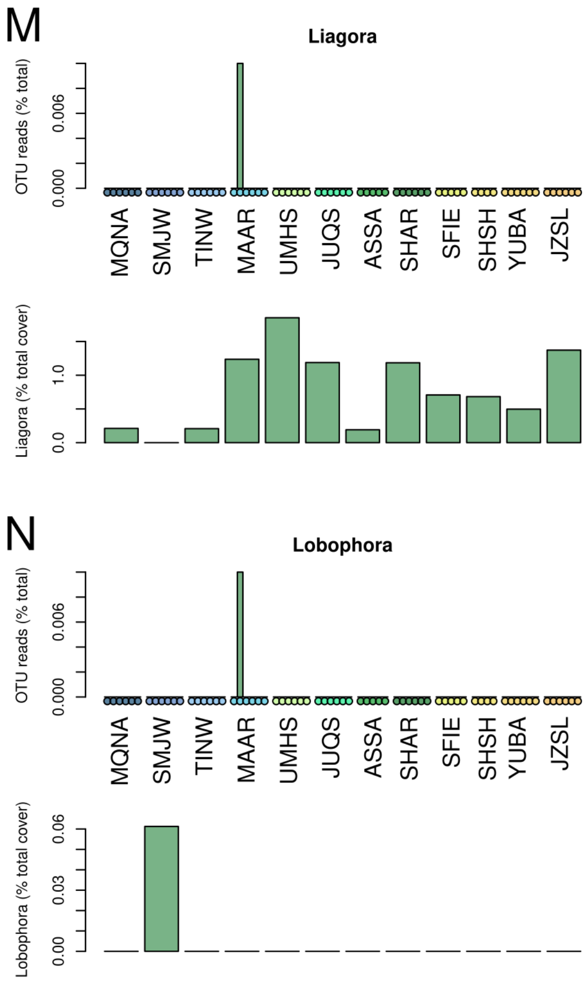


**Supplementary Figure 5:** Read counts (%) of 14 taxa across all samples and sites, compared to percent cover of the same taxa detected through the visual analysis of the photoquadrats.

### Supplementary Tables

**Supplementary Table 1.** Benthic biotic and abiotic categories (total 83) grouped into broader categories including Hard coral, dead hard coral, soft coral, invertebrates, algae, CCA, turf, bare rock, rubble, sand, tape and shadow

| **Benthic group** | **Benthic category** |
|---|---|
| **Hard coral** | *Acropora* branching |
| | *Acropora* tabular |
| | *Astreopora* |
| | Black Coral |
| | *Coscinarea* |
| | *Cyphastrea* |
| | *Dipsastrea* |
| | *Echinopora* encrusting |
| | *Echinopora* branching |
| | *Echinophyllia* |
| | *Favites* |
| | *Galaxea* |
| | *Gardineroseris* |
| | *Goniastrea* |
| | *Goniopora* |
| | Hard coral branching |
| | Hard coral columnar |

| | |
|---|---|
| | Hard coral encrusting |
| | Hard coral foliose |
| | Hard coral free living |
| | Hard coral massive |
| | Hard coral unknown |
| | *Leptastrea* |
| | *Leptoseris* |
| | *Lobophora* |
| | *Lobophyllia* |
| | *Merulina* |
| | *Millepora* |
| | *Montipora* encrusting |
| | *Montipora* massive |
| | *Oulophyllia* |
| | *Oxypora* |
| | *Pachyseris* |
| | *Paramontastrea* |
| | *Pavona* |
| | *Platygyra* |
| | *Plerogyra* |
| | *Pocillopora* |
| | *Podobacia* |
| | *Porites* columnar |
| | *Porites* encrusting |

| | |
|---|---|
| | *Porites* massive |
| | *Psammocora* |
| | *Siderastrea* |
| | *Stylocoeniella* |
| | *Stylophora* |
| | *Trachyphyllia* |
| | *Turbinaria* coral |
| **Dead hard coral** | Dead hard coral |
| **Soft coral** | *Melithea* |
| | Nephtheidae |
| | *Rhytisma* |
| | *Sarcophyton* |
| | *Sclerophytum* |
| | Unclassified soft coral |
| | *Tubipora* |
| | *Xeniidae* |
| **Invertebrates** | Bivalve |
| | *Tridacna* |
| | Sponge encrusting |
| | Sponge massive |
| | Unclassified invertebrate |
| | Zoanthid |
| **Algae** | Cyanobacteria |
| | *Dictyota* |

| | |
|---|---|
| | Encrusting macroalgae |
| | *Halimeda* |
| | *Liagora* |
| | Macroalgae green |
| | Macroalgae red |
| | Macroalgae |
| | *Turbinaria* algae |
| **CCA** | CCA |
| **Bare rock** | Bare rock |
| **Rubble** | Rubble |
| **Sand** | Sand |
| **Turf** | Turf dead coral |
| | Turf rock |
| **Other** | Debris |
| | Mobile fauna |
| | Unknown |
| **Tape** | Tape |
| **Shadow** | Shadow |

**Supplementary Table 3.** Site ID, Coordinates and distance to shore

| Site ID | Latitude | Longitude | Distance to shore (Km) |
|---|---|---|---|
| MAAR | 28.030893 | 34.574938 | 3.27 |
| SHAR | 27.619334 | 35.515665 | 0.01 |
| SHSH | 27.927334 | 34.898559 | 15.26 |
| SFIE | 27.924458 | 34.775869 | 13.71 |
| MQNA | 28.394166 | 34.736432 | 0.01 |
| SMJW | 28.177289 | 34.633719 | 0.02 |
| JUQS | 27.921689 | 35.239827 | 3.43 |
| YUBA | 27.774779 | 35.139886 | 20.73 |
| JZSL | 27.637252 | 35.302841 | 17.36 |
| TINW | 28.032631 | 34.51739 | 8.25 |
| UMHS | 28.048371 | 34.767467 | 1.00 |
| ASSA | 27.838777 | 35.265896 | 6.74 |

**Supplementary Table 4.** Validation of *Tridacna maxima* detection in photoquadrat surveys at sites JUQS, SMJW and UMHS.

| Site | Transect # | Number of images with manually detected Tridacna | Number of Tridacna indviduals manualy detected | Number of images with Tridacnas detection based on annotation points | Number of Tridacna detections based on annotation points |
|---|---|---|---|---|---|
| JUQS | 1 | 8 | 12 | 1 | 1 |
| JUQS | 2 | 11 | 17 | 6 | 7 |
| JUQS | 3 | 7 | 16 | 2 | 2 |
| SMJW | 1 | 0 | 0 | 0 | 0 |
| SMJW | 2 | 0 | 0 | 0 | 0 |
| SMJW | 3 | 1 | 1 | 1 | 1 |
| UMHS | 1 | 20 | 79 | 6 | 10 |
| UMHS | 2 | 19 | 105 | 5 | 12 |
| UMHS | 3 | 18 | 57 | 5 | 6 |
